# *Chlamydia* plasmid-encoded protein Pgp2 is a replication initiator with a unique β-hairpin necessary for iteron-binding and plasmid replication

**DOI:** 10.1101/2024.11.14.623704

**Authors:** Danny Wan, Matthew Pan, Guangming Zhong, Huizhou Fan

## Abstract

The virulence plasmid of the obligate intracellular bacterium *Chlamydia* encodes eight proteins. Among these, Pgp3 is crucial for pathogenicity, and Pgp4 functions as a transcriptional regulator of both plasmid and chromosomal genes. The remaining proteins, Pgp1, Pgp5, Pgp6, Pgp7, and Pgp8, are predicted to play various roles in plasmid replication or maintenance based on their amino acid sequences. However, the function of Pgp2 remains unknown, even though it is required for transformation. In this study, we utilized AlphaFold to predict the 3-dimensional (3-D) structure of *C. trachomatis* Pgp2. Despite a lack of apparent sequence homology, the AlphaFold structure exhibited high similarity to experimentally determined structures of several plasmid replication initiators. Notably, Pgp2 features a unique β-hairpin motif near the DNA-binding domain, absent in other plasmid replication initiators with overall 3-D structures similar to Pgp2. This β-hairpin motif was also present in AlphaFold models of Pgp2s across all 13 *Chlamydia* species. To assess its significance, we engineered a plasmid lacking the 11 amino acids constituting the β-hairpin motif in *C. trachomatis* Pgp2. Although this deletion did not alter the overall structure of Pgp2, the mutated plasmid failed to transform plasmid-free *C. trachomatis*. These findings reveal that Pgp2 is a plasmid replication initiator, with the β-hairpin motif playing a critical role in binding to its cognate iteron sequences in the replication origin of the chlamydial plasmid.

## INTRODUCTION

*Chlamydia* is an obligate intracellular bacterium pathogenic to humans and animals (1). For example, *Chlamydia trachomatis* is the leading cause of bacterial sexually transmitted infections globally and of preventable blindness in certain under-developed regions (2, 3). *C. muridarum* serves as a model for studying chlamydial diseases and immune responses in mice (4, 5).

All *Chlamydia* species carry a conserved plasmid. Although previously referred to as a “cryptic plasmid,” it is more accurately described as a virulence plasmid since studies involving mouse and non-human primate models demonstrated that plasmid-free *Chlamydia* strains exhibit significantly reduced pathogenicity (4-6). This highlights the possibility of targeting the plasmid to mitigate chlamydial infections.

The chlamydial plasmid encodes eight open reading frames, producing proteins designated Pgp1-8. Pgp3 is a key virulence factor, playing a crucial role in the progression of chlamydial infections from the lower genital tract to the upper genital tract (7-9) and in the colonization of the gastrointestinal tract (10). Pgp4 is a transcriptional regulator of Pgp3 and various chromosomal genes (11-13).

Three of the remaining six proteins are implicated in plasmid replication based on their amino acid sequences, while two are involved in plasmid maintenance. Specifically, Pgp1 is predicted to function as a DnaB helicase (14). Pgp5 and Pgp6 are predicted to act as ParA and ParB, respectively, mediating the partitioning of daughter plasmids (15). Pgp7 and Pgp8, both with homology to recombinase subunits XerC and XerD, are believed to mediate the resolution of plasmid multimers (15). Transformation studies have shown that plasmids lacking Pgp1, Pgp6, and Pgp8 fail to transform plasmid-free *C. trachomatis* and *C. muridarum*, whereas those lacking Pgp5 and Pgp7 do form transformants (12, 16). These findings indicate that Pgp1 and Pgp6 are essential for plasmid replication, Pgp8 is necessary for plasmid maintenance, and Pgp5 and Pgp7 are non-essential for replication and maintenance, likely due to functional redundancy with chromosome-encoded proteins.

In contrast to all the other plasmid-encoded proteins, the function of Pgp2 remains elusive due to a lack of sequence homology with known proteins. In the aforementioned transformation studies, Pgp2-deficient plasmids also failed to transform *C. trachomatis* and *C. muridarum*, suggesting that Pgp2 is necessary for plasmid replication or maintenance (12, 16). In this study, we predicted the structure of Pgp2 using AlphaFold and identified it as a plasmid replication initiator through DALI protein 3-D structural homology search. Additionally, we discovered a unique β-hairpin motif in Pgp2 that is absent in other replication initiators but highly conserved across *Chlamydia* species. An engineered plasmid lacking the β-hairpin motif (Pgp2Δ108-118) failed to transform *C. trachomatis*, providing strong evidence that Pgp2 functions as a plasmid replication initiator, with the conserved β-hairpin motif playing an essential role in its activity.

## RESULTS AND DISCUSSION

### Prediction of Pgp2 as a replication initiator

To determine the function of Pgp2, we first employed AlphaFold (17) to predict the 3-D structure of *C. trachomatis* Pgp2 (Fig. 1). The AlphaFold prediction yielded an average Local Distance Difference Test (pLDDT) score of 90.24 out of a maximum possible score of 100, indicating a reliable structural model (17). We then identified structural homologs of Pgp2 using the DALI server (18), comparing the AlphaFold-predicted structure against a database of known protein structures. The DALI search revealed significant structural resemblances between the Pgp2 structure and experimentally determined structures of several known plasmid replication initiators, including the π initiator of the *E. coli* R6K plasmid (PDB accession number 2NRA) (19), RepE of the *E. coli* F plasmid (PDB accession number 8AAN) (20), and RctB of *Vibrio cholerae* chromosome II (PDB accession number 5TBF), with root mean square deviation (RMSD) values of 3.3 Å, 3.3 Å, and 4.6 Å, respectively (Table 1). Notably, the *C. trachomatis* Pgp2 had Z-scores above 10 for all three matches, a DALI metric indicating how much the alignment score of a query structure with a target structure deviates from the mean score expected by chance. While a Z-score greater than 2 is generally considered significant for structural similarity, Z-scores above 8–10 represent highly similar and biologically relevant matches. These high Z-scores support the hypothesis that Pgp2 functions as a plasmid replication initiator, binding to its cognate DNA-binding site (iteron) sequences within the replication origin and recruiting the DNA helicase needed to initiate plasmid replication. The organization of Pgp1 (helicase) and Pgp2 as an operon further supports this Pgp2 functional prediction for Pgp2.

**Table 1.** Proteins with structural homology to the AlphaFold model of *C. trachomatis* Pgp2 identified by DALI search. The Z-score is an optimized similarity score defined as the sum of equivalent residue-wise Cα-Cα distances among two proteins. Abbreviations: RMSD, Root-mean-square deviation of atomic positions; lali, length of alignment in two proteins; nres, the number of residues in the matched structure.

| Rank | PDB accession # (reference) | Protein | Bacterium | Z-score | RMSD (Å) | lali | nres | Sequence identity (%) |
| --- | --- | --- | --- | --- | --- | --- | --- | --- |
| 1 | 2NRA (19) | $\pi$ Initiator | *E. coli* | 13.1 | 3.3 | 213 | 252 | 10 |
| 2 | 8AAN (20) | RepE | *E. coli* | 12.7 | 3.3 | 205 | 228 | 8 |
| 3 | 5TBF | RctB | *V. cholerae* | 10.8 | 4.6 | 209 | 280 | 10 |

**Fig. 1.**
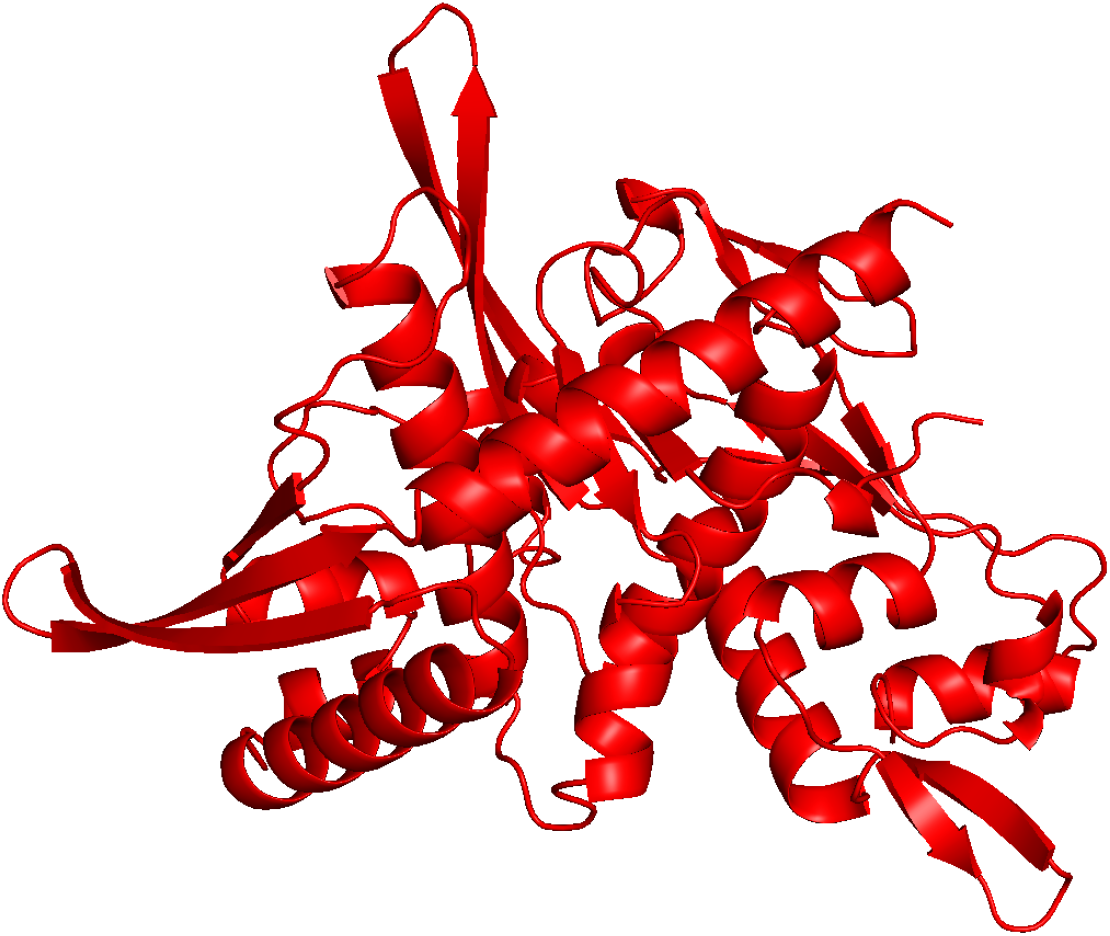
Three-dimensional structure of *C. trachomatis* L2 Pgp2. AlphaFold3 predicted the structure.

### Identification of a conserved β-hairpin motif in Pgp2 of *Chlamydia* species

Superimposition of the AlphaFold structure of *C. trachomatis* Pgp2 onto 2NRA, 8AAN, and 5TBF revealed a potentially distinct β-hairpin motif found only in Pgp2 but absent in the other initiators (Fig. 2A-C). The β-hairpin motif in Pgp2 extends from a helix whose equivalents in 2NRA and 8AAN bind their respective iterons (Fig. 2D, E). The 11 residues comprising this motif form two short β-strands (R108-Y109-K110 and R113-N114-K115-Y116-E117-F118) connected by a 2-residue loop (T111-S112) (Fig. 3A). Sequence alignment revealed that the residues constituting the β-hairpin motif in *C. trachomatis* Pgp2 are highly conserved across all 13 Chlamydia species (Fig. 3A); the same β-hairpin motif is present in the AlphaFold structures of all Pgp2s (Fig. 3B).

**Fig. 2.**
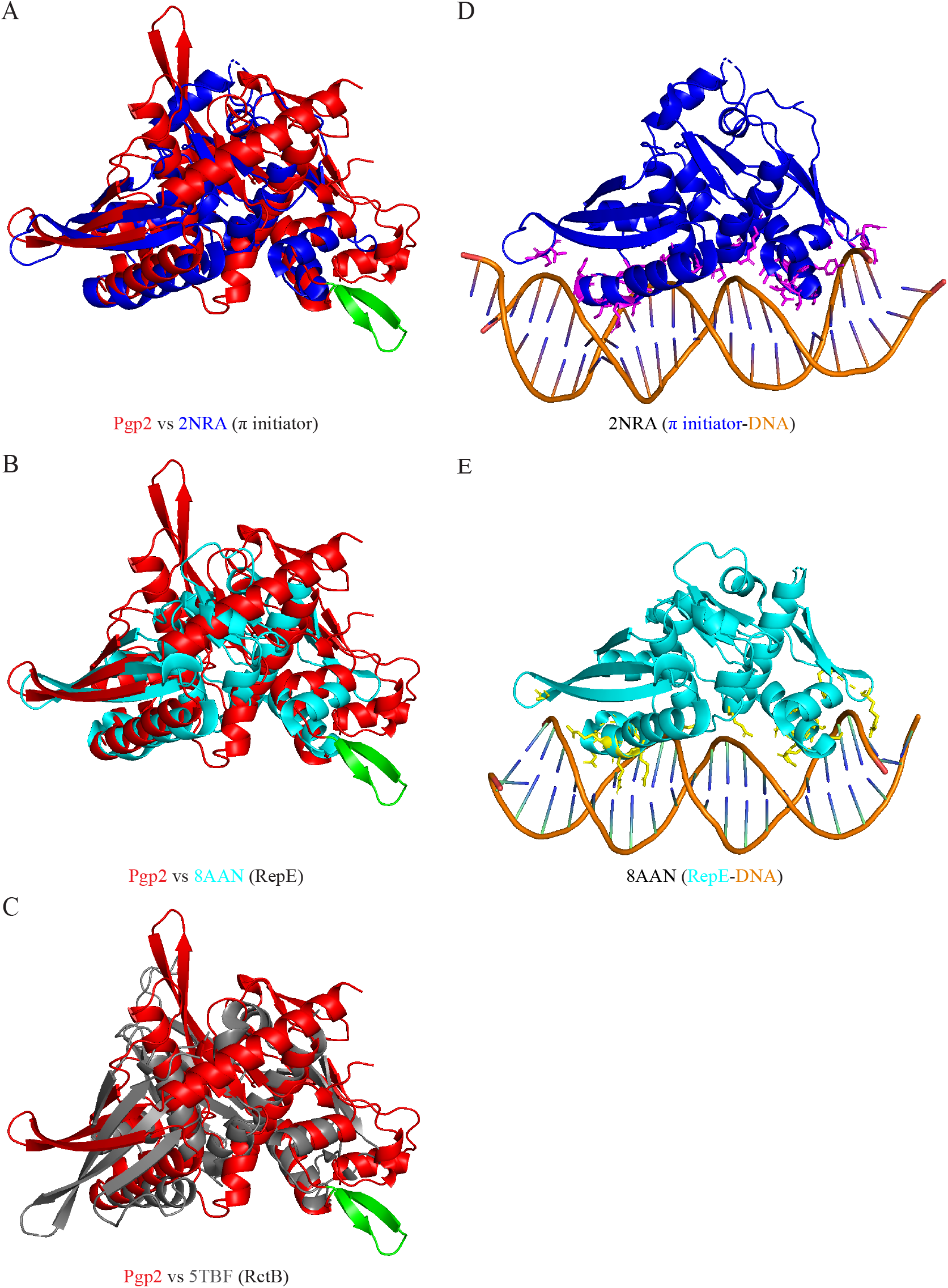
Identification of a β-hairpin motif extended from a helix whose equivalent in other replication initiators interacting with the iteron DNA. (A-C) Superimposition of AlphaFold-predicted Pgp2 structure (red) on the experimental structures of π initiator of the *E. coli* R6K plasmid (PDB accession number 2NRA) (blue), the RepE initiator of *E. coli* F plasmid (PDB accession number 8AAN) (teal), and the RctB initiator of *V. cholerae* (PDB accession number 5TBF) (gray). Pgp2 features an 11-residue β-hairpin (green) not seen in other replication initiators. (D, E) Experimental structure of 2NRA and 8AAN containing respective iteron DNA. Note that a comparison of panels D and E with A and B, respectively, reveals that the unique β-hairpin in Pgp2 is an extension of a DNA-binding helix.

**Fig. 3.**
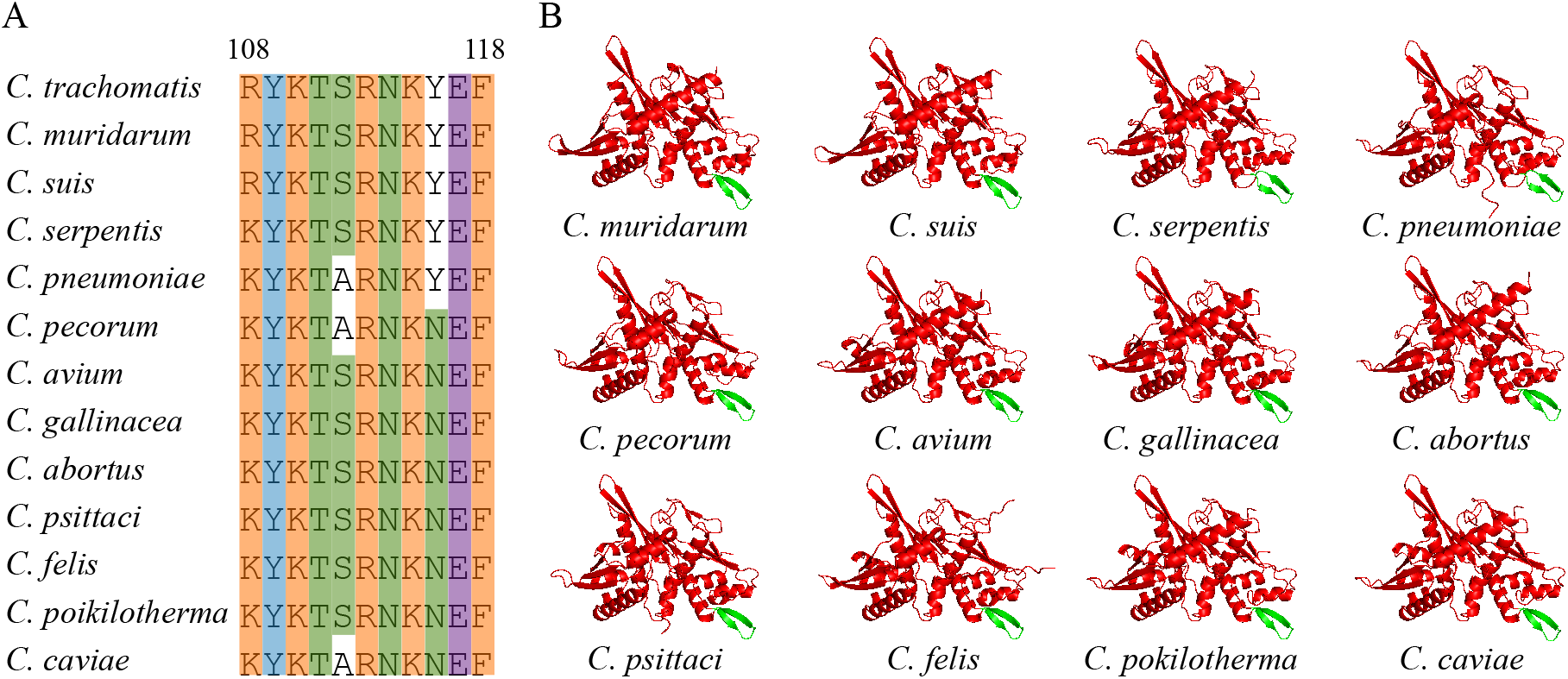
Conservation of the β-hairpin motif in Pgp2s of all *Chlamydia* species. (A) The amino acids constituting the β-hairpin in *C. trachomatis* Pgp2 are conserved in all 12 other species. Alignment of Pgp2s of all 13 *Chlamydia* species was performed using Clustal Omega, although only the 11 amino acid sequences that form the β-hairpin are presented. (B) Pgp2s of all 12 other *Chlamydia* species share the β-hairpin structure (green) with *C. trachomatis*. All structures were predicted using AlphaFold3.

### Requirement of the β-hairpin motif for Pgp2 activity

Given the conservation of the distinct β-hairpin motif identified through structural alignment near the predicted DNA-binding region, we hypothesized that the β-hairpin motif is required for Pgp2’s activity. To test this hypothesis, we removed the 33 nucleotides coding for R108-F118 of Pgp2 from pGFP::SW2, a shuttle vector containing the full sequence of the *C. trachomatis* L2 plasmid and a green fluorescence protein (GFP) gene for easy detection of transformants (Fig. 4A, B) (21). Other than the absence of the β-hairpin motif, the AlphaFold structure of the modified Pgp2 (Pgp2Δ111-118) was fully superimposable with the wild-type Pgp2 structure, indicating that removing the β-hairpin did not affect the overall structure (Fig. 4C).

**Fig. 4.**
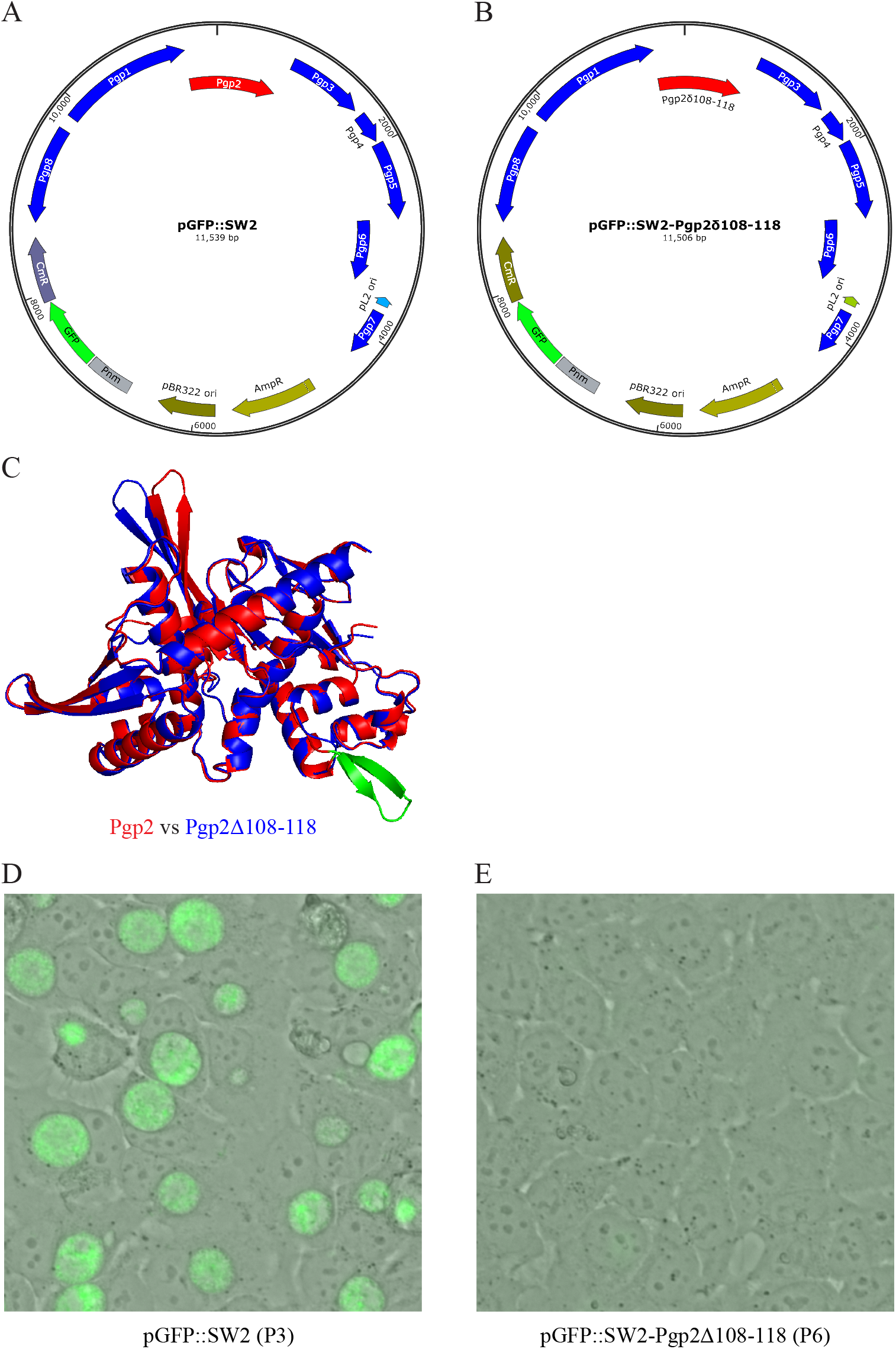
The distinct β-hairpin motif is required for Pgp2 activity. (A) The map of the shuttle vector pGFP::SW2 encoding wildtype Pgp2 (21). (B) The map of the shuttle vector pGFP::SW2-Pgp2Δ108-118 encoding the Pgp2 lacking residues 108-118 that form the unique β-hairpin. (C) Superimposition of wildtype Pgp2 and Pgp2Δ108-118. (D) A representative microscopic image of green fluorescence protein-expressing chlamydial inclusions formed in host cells in a passage 2 culture of *C. trachomatis* transformed with pGFP::SW2 and selected with ampicillin. (E) A representative microscopic image lacking fluorescence protein-expressing inclusions in a passage 6 culture of *C. trachomatis* transformed with pGFP::SW2-Pgp2Δ108-118 and selected with ampicillin. (D, E) Images were acquired at 36 hpi.

We transformed the plasmid-free *C. trachomatis* L2r strain (22) with pGFP::SW2 and pGFP::SW2-pgp2Δ111-118. Transformants were selected by passaging cultures in a medium containing ampicillin as the selection agent. In all three independent transformation experiments, pGFP::SW2 transformants with green fluorescence were detectable starting at passage 2. In contrast, no pGFP::SW2-pgp2Δ108-118 transformants were detected even by passage 5. Representative microscopic images of the pGFP::SW2 transformation culture with GFP-expressing chlamydiae at passage 3 and the pGFP::SW2-pgp2Δ108-118 transformation culture lacking GFP signals at passage 6 are shown in Fig. 4D and Fig. 4E, respectively. These results suggest that the conserved β-hairpin motif is necessary for Pgp2’s function. Most likely, the β-hairpin motif contributes to DNA binding. Although the helix-turn-helix motif is more commonly associated with DNA binding, β-hairpin motifs have also been shown to bind DNA in some cases [e.g., (23, 24)]. Notably, 4 of the 11 amino acids in the β-hairpin are positively arginine and lysine residues (Fig. 3A); they likely facilitate interactions with the negatively charged iteron DNA, contributing to Pgp2’s binding specificity and function.

In conclusion, this study reveals that Pgp2, the sole function-unknown protein encoded by the virulence chlamydial plasmid, is a DNA replication initiator. We have demonstrated the requirement of a conserved β-hairpin motif for Pgp2’s function, highlighting the possibility of targeting this structure as a strategy to disrupt plasmid-mediated virulence.

## MATERIALS AND METHODS

### Three-dimensional structure prediction

The amino acid sequences of Pgp2 from *Chlamydia* species, retrieved from the National Center for Biotechnology Information (NCBI) database, were used as inputs for structural prediction using the online AlphaFold 3 servers (17). The top-ranked predicted structures were selected for further analysis.

### 3-D-structure similarity analysis

The top-ranked AlphaFold-predicted structure of Pgp2 was submitted to the DALI server, and the resulting top hits were selected for superimposition. The superimposition was performed using PyMOL. The structural alignments were visually inspected to identify common structural motifs and unique features.

### Construction of vector expressing Pgp2 lacking the β-hairpin

pGFP::SW2-pgp2Δ108-118 was constructed by assembling two overlapping DNA fragments amplified using the shuttle vector GFP::SW2 carrying all the 8 Pgp genes (21) as the template. Fragment 1 was amplified using the forwarding primer named pgp2_cut_F1-F with the sequence 5’-gcttatggagttaagAGTGGAAAAGAAGCTGAAACT-3’ (the lowercase letters represent nucleotides coding for amino acids A102-K107 and the uppercase letters represent nucleotides coding for S119-T125) and the reverse primer US-pUC ori-F with the sequence 5’-gggattttggtcatgagattatc-3’ located in a noncoding sequence between the beta-lactamase open reading frame and the pBR322 replication origin. Fragment 2 was amplified using the forwarding primer US-pUC ori-R with the 5’-GATAATCTCATGACCAAAATCCC-3’, which is reverse complementary to US-pUC ori-F, and reverse primer pgp2_cut_F2-R with the sequence 5’-CTTTTCCactcttaactccataagcctctaaga-3’ (the lowercase letters represent nucleotides that is reverse and complementary to codons 100-117 and uppercases represent nucleotides that is reverse and complementary to codons 118 and 119 and part of codon 120). The NEB Q5 high-fidelity DNA polymerase (New England Biolabs) was used to amplify the overlapping fragments, which were assembled using the NEBuilder HiFi DNA Assembly kit (New England Biolabs) for the assembly. NEB 10-beta competent *E. coli* was transformed with the assembly product. Transformed colonies were subject to PCR-screening using diagnostic primers pgp2_cut_diag-F (5-ggagttaagAGTGGAAAA-3’) and pgp2_cut_diag-R (5’-taatcacccagtcgataaat-3’). Plasmid DNA from two colonies with positive amplification of the 391-bp diagnostic fragment was prepared and was subject to Nanopore whole plasmid sequencing at Quintara Biosciences (Boston). The plasmid with the confirmed deletion of codons 108-118 and without any additional mutations elsewhere was used to transform *Chlamydia*.

### *C. trachomatis* transformation

Transformation of the plasmid-free *C. trachomatis* variant L2r (22) with the parental shuttle vector pGFP::SW2 or its derivative pGFP::SW2-pgp2Δ108-118 was performed as described (25) with modifications (26). Briefly, 10^7^ inclusion-forming units of EBs were mixed with 5 μg of plasmid DNA in 50 μl CaCl2 buffer (10 mM Tris, pH 7.4 and 50 mM CaCl2) and incubated for 30 min at room temperature. The mixture was then diluted with 12 mL Hanks Balanced Salt Solution and used to inoculate a 6-well plate of confluent L929 cells (i.e., ∼2 ml of the suspension per well). Monolayers were infected at room temperature by centrifugation for 1 h at 900×g, after which HBSS was replaced with DMEM containing 5% FBS and 1 μg/mL cycloheximide (2 ml/well). Ampicillin (sodium salt) was added to the cultures at 2 hpi to achieve the final 10 µg/mL concentration. The passage 0 cultures were harvested at 36 hpi and passaged onto new 6-w plates at a 1-to-1 passage ratio using centrifugation-assisted procedures described above. Ampicillin was added to the cultures at the time of inoculation. The cultures were passaged every other day until green fluorescence protein-expressing inclusions were detected under an epifluorescence microscope. Since transformed chlamydiae are typically observed in P1 to P3 cultures in our lab, transformations were considered unsuccessful if GFP-expressing inclusions were not detected in P5 cultures.

### Imaging of *C. trachomatis* in host cells

HeLa229 cells grown on 6-well plates were inoculated with harvests of a P2 culture of the pGFP::SW2 transformation or of a P5 culture of the pGFP::SW2Δ108-118. At 36 hpi, the culture medium was replaced with phosphate-buffered saline. Images were acquired on an Infinity i8-3 CMOS monochrome camera. Image pseudo-coloring and overlay were performed using the ACINST03 software (27).

## ACKNOWLEDGMENTS

We thank Lingling Wang for performing the transformation experiments and Yuxuan Wang for acquiring microscopic images. This work was supported by the National Institutes of Health grant # AI154305 (to HF) and AI182210 (to GZ and HF).

